# A flexible diet platform for the nutrigenomic screening of *Drosophila* disease models

**DOI:** 10.64898/2026.03.01.708389

**Authors:** Zoriana Novosiadla, Sarah Mele, Emily Kerton, John Christodoulou, Sebastian Dworkin, Matthew D. W. Piper, Travis K. Johnson

## Abstract

Nutrient-gene interactions shape metabolic disease phenotypes, but systematic testing is limited by the effort required to produce defined diets. We developed a flexible protocol that assembles precisely defined synthetic diets for *Drosophila melanogaster* from individual stock solutions. Using this approach, we generated a rational array of 51 single-nutrient-varied diets and demonstrated that flexible and standard preparation methods produce comparable developmental timing, survival, adult body weight, and starvation resistance in wildtype flies. Applying the dietary array to a *Drosophila* model of isolated sulfite oxidase deficiency, a severe condition caused by impaired sulfur amino acid catabolism, revealed nutrient-specific effects on pupal survival and pupariation timing, including rescue by cysteine depletion and other amino acid modified diets. This platform provides a scalable *in vivo* framework for mapping genotype-specific nutritional responses across *Drosophila* disease models.

**Key points:**

- Developed a flexible platform for assembling precisely defined synthetic diets in *Drosophila*.
- Generated a rational 51-diet array, enabling scalable nutrigenomic screening.
- Flexible and standard diet preparations yielded matching developmental and adult fitness outcomes.
- Dietary array screening in a sulfite oxidase deficiency model recapitulated cysteine sensitivity and identified novel nutrient modifiers.

## Introduction

Dietary composition is a major environmental determinant of phenotype, influencing health, fitness and disease susceptibility across taxa (Holmes et al., 2008; Meddens et al., 2021). Nutrient-gene interactions are particularly relevant in metabolic disorders, where altered enzymatic or transport capacities modulate how organisms respond to specific dietary components (DeBerardinis and Thompson, 2012; Vander Heiden and DeBerardinis, 2017). Despite their importance, these interactions remain challenging to investigate systematically because of the ultra-low prevalence of metabolic disorders in the population (Waters et al., 2018). The use of traditional model systems that capture the organismal complexity of these disorders, such as rodents, has provided valuable insights into nutrient-disease interactions (Zinnanti et al., 2009; Vogel et al., 2014). However, the testing of many varied diets on these animals remains time-consuming and costly, and the diets themselves can be difficult to source and variably defined, or challenging to manipulate at scale.

More recently, the development of defined and semi-defined diets for *Drosophila*, has enabled mechanistic studies spanning basic biology and disease modelling (Piper et al., 2014; Lesperance and Broderick, 2020; Martelli et al., 2024a; Sorge et al., 2025). While these diets open the door to detailed exploration of nutrient-gene interactions in an animal model, the generation of highly varied diet arrays remains limited by labour-intensive and time-consuming preparation steps. This is because each of the 40+ individual ingredients must be weighed, solubilised and batched in a reproducible and scalable manner. This constraint is particularly relevant for amino acids, as their precise stoichiometric balance can impact protein synthesis efficiency and quality (Millward et al., 1988; Harper et al., 1993).

*Drosophila* is a powerful system for disease modelling, with >75% of human disease-causing genes having functional fly orthologs (Yamamoto et al., 2014). This extends to inherited metabolic disorders (IMDs), a group of more than 1,400 distinct conditions that are known for their response to therapeutic dietary manipulation (Ferreira et al., 2021; Mele et al., 2023) *Drosophila* possess the same core pathways of amino acid metabolism, mitochondrial function and redox homeostasis as humans (Adams et al., 2000; Leopold and Perrimon, 2007). There are several *Drosophila* models of IMDs including organic acidemias, sulfur-metabolism disorders, transporter deficiencies and fatty-acid oxidation disorders, studies of which have provided functional and therapeutic insights (Bellen et al., 2010; Wangler et al., 2017; Martelli et al., 2024b). Since IMDs, specifically amino acid disorders, are frequently diet-responsive, and trials of novel nutrient combinations in the clinic are restricted by the low prevalence of IMDs, the *Drosophila* model is emerging as a useful platform for nutrigenomic screening (Bangi et al., 2021; Martelli et al., 2024b; Mele et al., 2025).

To address current limitations in dietary manipulation, we developed a flexible diet preparation workflow that assembles precisely defined synthetic diets by combining individual nutrient solutions with a Holidic diet base formula. In principle, this approach can generate arbitrary nutrient compositions, enabling rapid construction of large arrays (feasibly tens to hundreds of diets) either manually or using semi-automated liquid handling robotics. Here, we show that this approach recapitulates phenotypes equivalent to those obtained with the standard preparation protocol. We then demonstrate its versatility by generating and testing a 51-diet array to map nutrient-dependent survival and developmental responses in a *Drosophila* model of isolated sulfite oxidase deficiency. This work establishes a scalable *in vivo* framework for systematic nutrigenomic investigations in *Drosophila*.

## Results and Discussion

### Establishment and validation of a flexible diet assembly platform

The Holidic diet is prepared by combining 46 individual purified nutrient components, each either weighed separately or added from stocks through multiple steps (Piper et al., 2014; Martelli et al., 2024). Under the standard workflow, most diet combinations are prepared independently, making it time-, and labour-intensive to produce multiple nutrient-varied diets. To enable scalable high-throughput dietary screening, we developed a flexible protocol in which a Holidic base medium containing all nutrients except amino acids was prepared in bulk. Defined amino acid mixtures were then assembled from individual amino acid solutions using a liquid-handling robot and then combined with the base medium to generate the final experimental diets **(Figure 1A)**. This approach allows individual amino acids to be varied systematically; however, it could also be used to vary other nutrients (*e*.*g*. vitamins, trace minerals) with an appropriate base media that lacks these components. To verify the performance of the new workflow, we reared wildtype *Drosophila* on media prepared using the flexible assembly protocol and assessed outcomes relative to the standard preparation method (as described in Martelli et al., 2024). No differences in survival nor developmental timing to the pupal stage, were observed between media preparation methods (**Figure 1B-D**). We also saw no differences between the adult weights attained on the two diets, suggesting no negative impact on growth (**Figure 1E**). Their ability to withstand starvation as adults was also not affected by diet preparation method, suggesting equivalent nutritional provisioning (**Figure 1F**). The reproducibility of these phenotypes validates the use of a scalable synthetic media preparation system.

**Figure 1.**
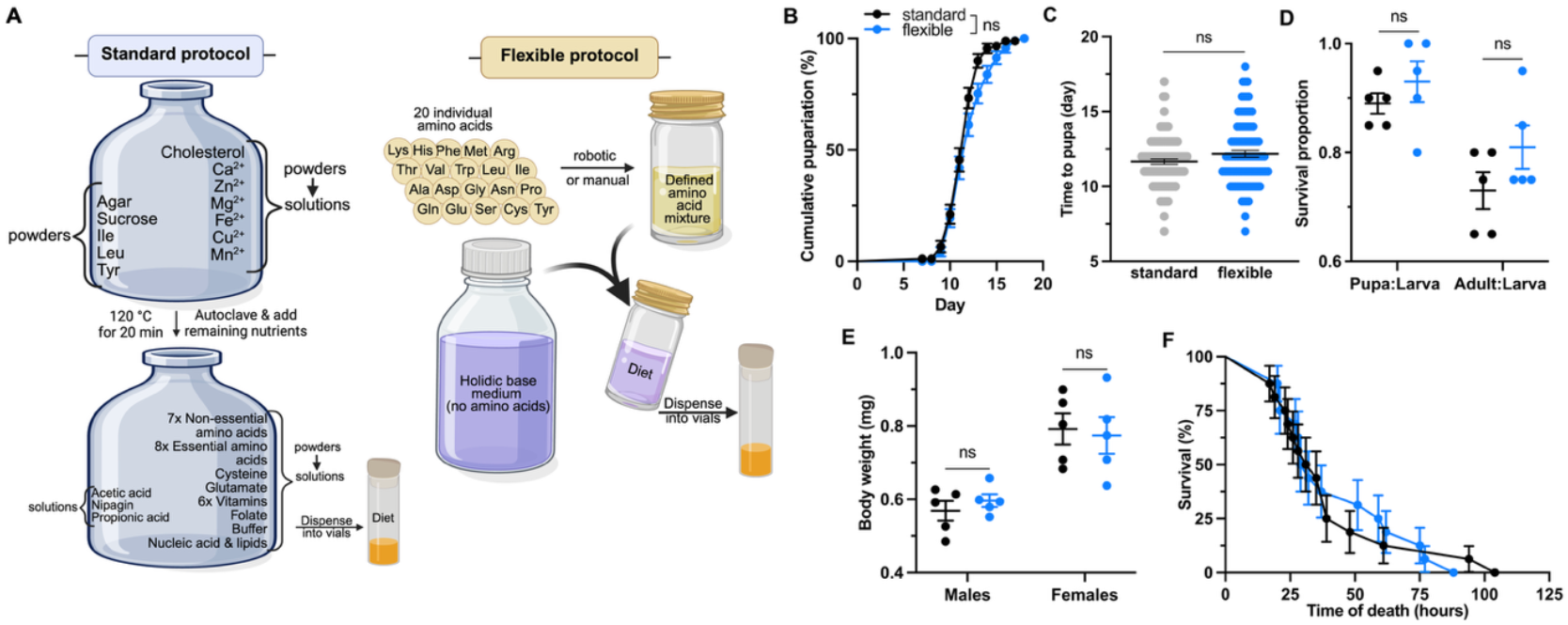
Development and validation of a flexible Holidic diet preparation workflow. **(A)** Schematic comparison of standard and flexible diet preparation protocols. In the standard protocol, 46 individual components are sequentially combined through multiple preparation steps, with some weighed and added directly during cooking and others added from stock solutions prepared from powdered reagents. In the flexible protocol, a Holidic base medium containing all nutrients except amino acids (AA) is prepared in bulk, while individual AA stock solutions are used to generate defined amino acid mixtures, which can be performed manually or using liquid-handling robotics. The mixtures and base medium are then combined to generate experimental diets, which are dispensed into vials. Comparison of developmental timing **(B, C)**, survival **(D)**, adult body weight **(E)** and adult male starvation resistance **(F)** of wild-type flies reared on diets prepared using standard (black) and flexible (blue) protocols. (**B-E**) Five replicates (vials) per diet, **(B-D)** n=100 larvae per diet. (**F**) n=16 adult males per diet. Cumulative pupariation and survival curves were analysed using the log-rank (Mantel– Cox) test, and median event times were calculated. Differences in pupariation timing and viability between groups were analysed using the Mann–Whitney U test. Body weight was compared using an unpaired two-tailed *t*-test. Error bars represented as mean ± 1 S.E. (ns, not significant). Created in BioRender. Novosiadla, Z. (2026). https://BioRender.com/iw408iv

Given our particular interest in IMDs affecting amino acid metabolism and their well-known responses to dietary manipulations (Woolf et al., 1955; Martelli et al., 2024; Mele et al., 2025), we designed a dietary array that systematically varies individual amino acid concentrations (**Table S1** and **S2**). Diets completely lacking essential amino acids (EAAs) were excluded from the array, as they are indispensable for protein synthesis and tissue maintenance, and required for sustaining life (Koehnle et al., 2004). Instead, EAAs were reduced to either 50% or 75% of the complete formula based on their effects observed on wildtype flies (Martelli et al., 2024). Non-essential amino acids (NEAAs) were reduced to 0% and 50% relative to the complete diet, and all amino acids were tested in excess at 150%. In total, this yielded 51 distinct diets devised to serve as a first-pass screen for *Drosophila* IMD models. Alternative designs could be used to test other nutrient classes, perform dose-response analysis, or vary two or more nutrients in combination.

### Systematic nutrigenomic profiling in a Drosophila model of sulfite oxidase deficiency

We applied the nutrient array to a *Drosophila* model of isolated sulfite oxidase deficiency (*shop*^*C15*^; Otto et al., 2018) to test whether our dietary array could uncover nutrient-disease interactions beyond those already known from targeted studies (Martelli et al., 2024). The *shop*^*C15*^ model represents an impairment in sulfur amino acid (methionine, Met, and cysteine, Cys) metabolism that closely mirrors the human condition, which is characterised by accumulation of neurotoxic intermediates such as sulfite, thiosulfate and S-sulfocysteine, resulting in rapid neurodegeneration and lethality (**Figure 2A**; Grings et al., 2013). In this pathway, Met serves as the precursor for Cys biosynthesis, with serine (Ser) providing the carbon backbone *via* the trans-sulfuration pathway (Finkelstein, 2000). Previous studies showed that restricting Met, Ser or Cys individually or in combination rescues lethality of *shop*^*C15*^ to varying degrees, with the strongest effect observed upon Cys depletion (Martelli et al., 2024).

**Figure 2.**
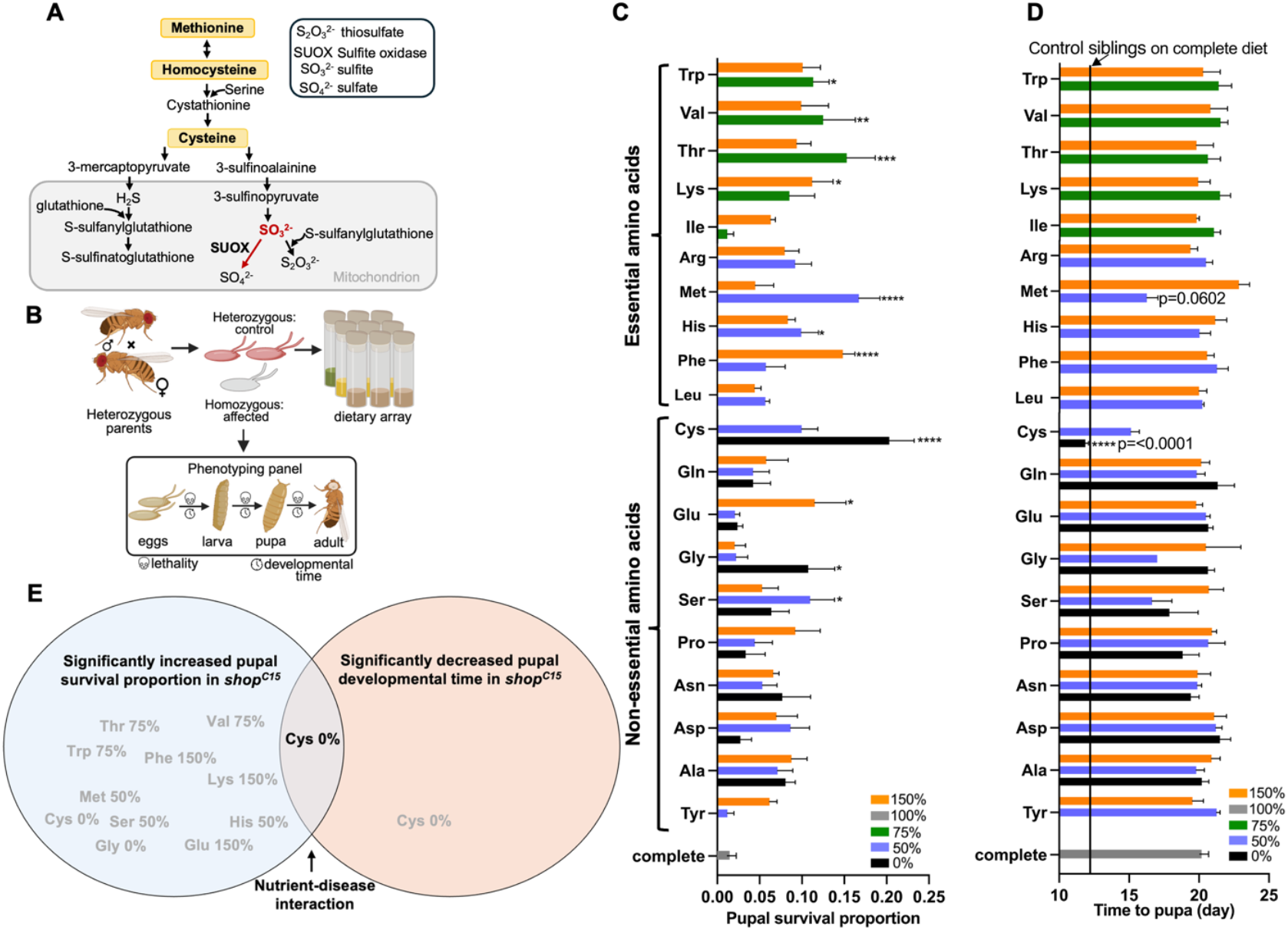
Nutrient-phenotype interactions revealed by amino acid modulation in the *Drosophila shop*^*C15*^ model. **(A)** Schematic of methionine (Met)-cysteine (Cys) metabolic pathway, highlighting key intermediates and the site of sulfite oxidase (SUOX) activity. **(B)** Schematic of the experimental design. Heterozygous *shop*^*C15*^*/FM7a* females were mated with *FM7a/Y males* to produce offspring: hemizygous (*shop*^*C15*^/Y) affected males (no fluorescent balancer) and control unaffected siblings (fluorescent balancer). Embryos were placed onto 51 diets (50 nutrient-varied and one complete diet, 5 replicates each) and monitored for 25 days. Survival of *shop*^*C15*^ hemizygotes was calculated as the proportion of total pupae. **(C)** *shop*^*C15*^ pupal survival proportions across diets varying essential and non-essential amino acids relative to the control diet. **(D)** *shop*^*C15*^ pupariation time across diets varying essential and non-essential amino acids relative to the control diet. The vertical line indicates the mean pupariation time of control siblings reared on the complete diet. **(E)** Venn diagram summarising significant nutrient-disease interactions. Error bars represent mean ± 1 S. E. One-way ANOVA with multiple comparisons (Tukey’s HSD) (\*\*\*\**P*<0.0001, \*\*\**P*<0.001, \*\**P*<0.01, \**P*<0.05).

Consistent with these findings, we detected significantly improved survival of *shop*^*C15*^ individuals on the Cys-free diet, as well as upon restriction of Met and Ser (**Figure 2B-E**), thereby providing strong validation of our approach. Complete lethality was observed on the 150% Cys diet, suggesting these flies are hypersensitive to Cys levels (**Figure 2C**). Notably, these diets did not impact the survival of sibling controls (**Figure S1A**). Beyond these nutrient interactions, our diet array also revealed several distinct amino acid perturbations that significantly increased pupal survival of *shop*^*C15*^ mutants toward the expected Mendelian ratio of 0.25 (**Figure 2C**). This included moderate restrictions of the essential amino acids Thr, Val, Trp, and His. A shared feature of these amino acids is their capacity to engage amino acid stress-response pathways (*e*.*g*., GCN2 and mTOR; De Vito et al., 2018; Zhang et al., 2025) when limited in the diet, raising the possibility that such responses contribute to protection in *shop*^*C15*^ mutants. We also saw elevated survival upon Gly removal, which could impact Ser availability *via* its generation in the glycine cleavage system, thereby mirroring Ser restriction (Kikuchi et al., 2008). Supplementation of Phe, Lys and Glu likewise improved survival. One possibility is that these alter amino acid transport and nutrient handling rather than having direct pathway-specific effects. For example, Glu may improve *shop*^*C15*^ survival by influencing nitrogen balance and/or cystine (oxidised Cys-Cys) uptake *via* systems like the cystine-Glu exchanger, (Fujii et al., 2023; Koppula et al., 2018). Similarly, Phe and Lys may act through altered competition within neutral (LNAA-type, Pietz et al., 1999) and cationic/dibasic (*e*.*g*. System *B*^*0,+*^ -like) amino acid transport systems, respectively (Satsu et al., 1998; Ellingsen et al., 2021).

We also investigated whether these diets could normalise the pronounced developmental delay observed for *shop*^*C15*^ larvae (*shop*^*C15*^ pupariation occurs at ∼day 20 compared to controls at day 12; **Figure 2D**). Among all diets tested, only the Cys-free diet shifted developmental timing significantly towards control values (**Figure 2D**). Reduced Met trended towards improvement but this was not significant (p=0.060). The developmental timing of sibling controls was largely unaffected by the diets, indicating that the effects observed in *shop*^*c15*^ mutants are genotype-specific (**Figure S1B**). The only exception was for the 0% and 50% Tyr diets, which significantly delayed pupariation, as has been previously described (Kosakamoto et al., 2024). Together, these findings demonstrate that Cys depletion exerts a significant and highly nutrient-specific normalising effect on both survival and developmental timing phenotypes in *shop*^*C15*^ mutants. Previous clinical dietary management strategies for this disorder have typically reduced both Cys and Met concurrently (Touati et al., 2000). Our data suggest that Cys restriction alone may provide benefit and therefore warrants further investigation as a targeted translational dietary strategy. Further work is needed to uncover the mechanisms by which other diets improved *shop*^*c15*^ survival and determine whether these findings can be translated to other pre-clinical models.

The synthetic diet preparation protocol described here provides a readily accessible means of nutrigenomic exploration across several to hundreds of precisely defined and reproducible diets. This enables focused nutrient manipulations including dose-response and multidimensional nutrient analyses, to broad untargeted screens and novel interaction discovery. While we demonstrated its use here for a single monogenic rare disease model, our nutrigenomics array is directly transferable across *Drosophila* genetics as a systematic first-pass framework to chart diet-genotype-phenotype interactions. By doing so, it permits mechanistic dissection in nutritional genomics, and aids in precision nutrition to accelerate the discovery of rational dietary interventions for disorders with strong metabolic bases.

## Materials and Methods

### Drosophila lines and maintenance

The following fly stocks were obtained from the Bloomington (BL) *Drosophila* Stock Centre: *y*^*1*^, *w*^*1118*^ as the wild-type strain (BL6598); FM7a, ChFP (BL35522) and *shop*^*C15*^ (kind gift from Christian Klämbt). The *shop*^*C15*^ mutant allele was balanced over a fluorescently labelled balancer chromosome (FM7a, ChFP). Flies were maintained at 22°C in 12 h:12 h light:dark conditions, and experiments were performed at 25°C.

### Synthetic media preparation

The standard synthetic diet preparation refers to the methods described in Martelli et al., 2024 and Piper et al., 2014. The method was modified to enable the semi-automated generation of multi-nutrient arrays. Firstly, a synthetic base medium (Martelli et al., 2024) (1.5L) lacking all amino acids was made. Individual stock solutions for each amino acid were prepared in a final volume of 100 mL MilliQ water after adjusting pH 4.5 with HCl followed by filter sterilisation **(Table S1)**. Tubes were stored at 4°C (up to a maximum 2 months) and visually inspected for precipitation and vortexed before use. For the dietary array, the volume of the amino acid is indicated in the “Volume (mL) of individual amino acid stock required per 35 mL of Holidic base medium to prepare the diet” column in **Table S2**. Where nutrients are omitted or reduced in concentration, the volume is adjusted according with water. The defined amino acid mixtures for each of the 51 diets were added to tubes containing 35 mL of molten synthetic base medium and mixed using a motorized pipette. Diets were dispensed at 6 mL per vial.

### Developmental timing and viability

Wild-type embryos were collected from population cages that were allowed to lay for 8 h on apple juice agar supplemented with yeast paste. Embryos were collected into baskets and washed with distilled water and ∼50 µL of embryos transferred onto plates containing synthetic media using a micropipette and a tip cut to widen the bore. Plates were incubated for 24 h until hatching. Newly hatched first instar larvae were transferred to vials (20 larvae per vial) containing the appropriate diet using a blunt metal probe. Vials were checked every 24 h for the presence of pupae and adults for 20 days. Pupariation kinetics were visualized as the cumulative number of individuals at each developmental stage over time (days post-egg lay). Median developmental time and survival proportion (pupa per larva; adult per pupa) were calculated.

### Nutrigenomics on shop^C15^

Large numbers of developmentally synchronised *shop*^*C15*^/FM7a, ChFP embryos were obtained from multiple population cages containing approximately 200 females and 50 males. Embryos were transferred to a 1.7mL tube, resuspended in phosphate-buffered saline (PBS), and allowed to settle for 1 minute. Excess PBS was removed, then 3 µL of embryos were dispensed into vials using a micropipette onto the medium surface. This was performed for each of the 51 modified diets replicated 5 times. Vials were monitored daily for 25 days. Total numbers of pupae were counted, and the proportion of surviving *shop*^*c15*^ hemizygotes (*shop*^*C15*^/total pupae) was calculated.

### Adult body weight measurement

Newly eclosed adult male and female flies were weighed using a Mettler-Toledo XS3DU Excellence Microbalance. Flies were weighed in groups of 10 with five biological replicates per diet. Mean individual weight was estimated by dividing total group weight by 10.

### Starvation assay

Flies were reared from egg to adult on Holidic diets prepared using either the standard or a flexible protocol. On days 3–4 post-eclosion, 16 males per diet were collected, briefly anesthetized with CO_2_(<1 min), and placed individually into wells of a 48-well plate containing 600 µL of 1% (wt/vol) agar in water. Wells were sealed with PCR plate sealing film (SealPlate™, Excel Scientific Inc.) and 20 small ventilation holes per well were made using a 20-guage needle. Wells were imaged every hour for 5 days using a custom-made robotic imaging system with infrared illumination in darkness (T.K. Johnson, un-published). Image series from each well were collated in ImageJ and assessed for the hour at which each fly stopped moving (time of death).

### Software and data analysis

All statistical analyses and data visualizations were performed using GraphPad Prism version 10.0 (GraphPad Software).

## Acknowledgements

We thank the Bloomington *Drosophila* Stock Centre, and Professor Christian Klämbt (University of Münster, Germany) for fly stocks, the Australian *Drosophila* Biomedical Research Facility (Ozdros) for stock importation, and members of the Johnson, Piper and Dworkin labs for helpful discussions. John Christodoulou is generously supported by The Royal Children’s Hospital Foundation as The Chair in Genomic Medicine.

## Author contributions

Z.N. designed the methodology and experiments, performed experiments, collected and analysed the data, generated the figures and co-wrote the manuscript. S.M. co-produced the figures and edited the manuscript. E.K. assisted with experiments and edited the manuscript. J.C. edited the manuscript. S.D. edited the manuscript and co-supervised the work. T.K.J. and M.D.W.P. conceived the study, acquired funding, supervised the work and co-wrote the manuscript.

## Data and resource availability

All data and reagents generated in this study are available from the corresponding author upon reasonable request.

