## Supplementary figures and tables for "A flexible diet platform for the nutrigenomic screening of *Drosophila* disease models"

**Supplementary Data**

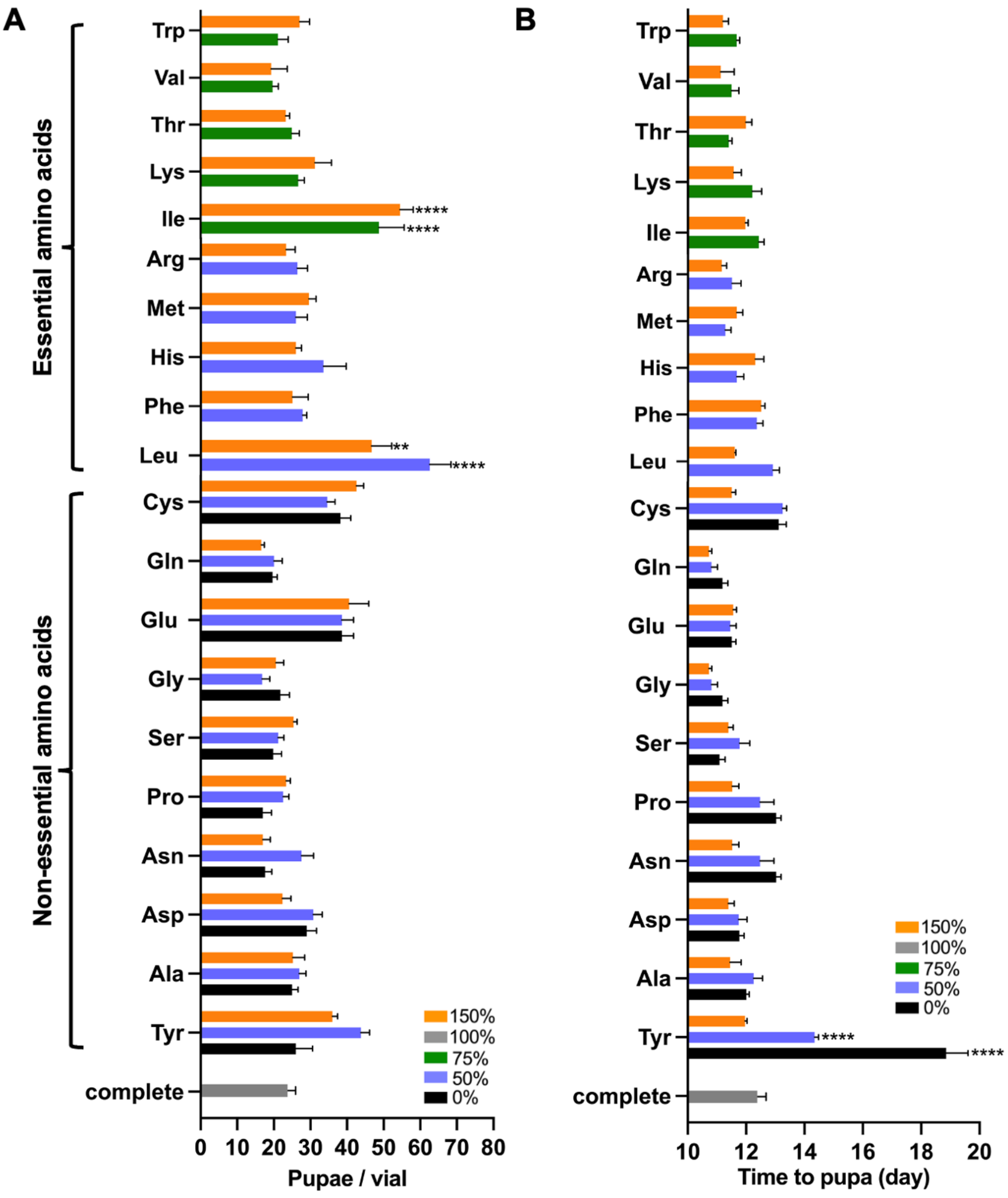


**Figure S1. Pupal counts and developmental timing of sibling controls of *shop*$^{C15}$ across amino acid-modified diets. (A)** Number of sibling control pupae per replicate vial. No diets negatively affected survival. **(B)** Developmental timing of sibling controls across diets. Data are presented as the mean ± 1 S. E. of 5 replicate vials. One-way ANOVA with multiple comparisons (Tukey's HSD, ****$P<0.0001$, **$P<0.01$).

**Table S1. Concentrations of amino acids used in the dietary array.**

| | | Concentration of individual stock solution | Final concentration of amino acid in the diet |
|---|---|---|---|
| | | g/100 ml of $H_2O$ | g/100 ml |
| **Essential amino acids** | **Phe** | 1.66 | 0.054 |
| | **His** | 1.08 | 0.035 |
| | **Ile** | 1.20 | 0.060 |
| | **Lys** | 2.25 | 0.074 |
| | **Leu *** | 2.20 | 0.110 |
| | **Met** | 1.00 | 0.032 |
| | **Arg** | 2.69 | 0.088 |
| | **Thr** | 1.83 | 0.060 |
| | **Val** | 1.98 | 0.065 |
| | **Trp** | 0.53 | 0.017 |
| **Non-essential amino acids** | **Ala** | 1.82 | 0.059 |
| | **Cys** | 0.36 | 0.018 |
| | **Asp** | 1.93 | 0.063 |
| | **Glu †** | 1.64 | 0.082 |
| | **Gly** | 1.27 | 0.041 |
| | **Asn** | 1.70 | 0.055 |
| | **Pro** | 1.61 | 0.053 |
| | **Gln** | 1.85 | 0.060 |
| | **Ser** | 2.27 | 0.074 |
| | **Tyr ‡** | 1 | 0.05 |

*Leu requires 500 µL of 37% HCl to dissolve in stock solution
†Glu (free acid) requires dropwise (~30 µL) of 2M NaOH to dissolve in stock solution
‡Tyr requires 750 µL of 37% HCl to dissolve in stock solution

**Table S2. Volumes of amino acid stock solutions used to generate the dietary array.**

| | | Volume (mL) of individual amino acid stock required per 35 mL of Holidic base medium to prepare the diet | | | |
|---|---|---|---|---|---|
| | | **50%** | **75%** | **100%** | **150%** |
| **Essential amino acids** | **Phe** | 0.568 | no diet | 1.135 | 1.703 |
| | **His** | 0.567 | no diet | 1.135 | 1.702 |
| | **Ile** | no diet | 1.313 | 1.750 | 1.750 |
| | **Lys** | no diet | 0.863 | 1.150 | 1.150 |
| | **Leu** | 0.875 | no diet | 1.750 | 2.625 |
| | **Met** | 0.562 | no diet | 1.125 | 1.687 |
| | **Arg** | 0.572 | no diet | 1.144 | 1.716 |
| | **Thr** | no diet | 0.863 | 1.150 | 1.150 |
| | **Val** | no diet | 0.861 | 1.148 | 1.148 |
| | **Trp** | no diet | 0.842 | 1.122 | 1.122 |
| **Non-essential amino acids** | **Ala** | 0.568 | no diet | 1.136 | 1.704 |
| | **Cys** | 0.875 | no diet | 1.750 | 2.625 |
| | **Asp** | 0.570 | no diet | 1.140 | 1.710 |
| | **Glu** | 0.875 | no diet | 1.750 | 2.625 |
| | **Gly** | 0.567 | no diet | 1.133 | 1.700 |
| | **Asn** | 0.566 | no diet | 1.133 | 1.699 |
| | **Pro** | 0.575 | no diet | 1.149 | 1.724 |
| | **Gln** | 0.568 | no diet | 1.135 | 1.703 |
| | **Ser** | 0.569 | no diet | 1.139 | 1.708 |
| | **Tyr** | 0.875 | no diet | 1.750 | 2.625 |

Ile, Lys, Thr, Val and Trp at 50% cause significant developmental delay in wildtype flies, these AAs were therefore tested at 75%.

The 51 diet array comprises the above 50%, 75%, and 150% variations, plus 10 non-essential amino acid 0% diets and the 100% control diet.
